# How many replicates to accurately estimate fish biodiversity using environmental DNA on coral reefs?

**DOI:** 10.1101/2021.05.26.445742

**Authors:** Salomé Stauffer, Meret Jucker, Thomas Keggin, Virginie Marques, Marco Andrello, Sandra Bessudo, Marie-Charlotte Cheutin, Giomar Helena Borrero-Pérez, Eilísh Richards, Tony Dejean, Régis Hocdé, Jean-Baptiste Juhel, Felipe Ladino, Tom B. Letessier, Nicolas Loiseau, Eva Maire, David Mouillot, Maria Mutis Martinezguerra, Stéphanie Manel, Andrea Polanco Fernández, Alice Valentini, Laure Velez, Camille Albouy, Loïc Pellissier, Conor Waldock

## Abstract

Quantifying the diversity of species in rich tropical marine environments remains challenging. Environmental DNA (eDNA) metabarcoding is a promising tool to face this challenge through the filtering, amplification, and sequencing of DNA traces from water samples. However, the reliability of biodiversity detection from eDNA samples can be low in marine environments because eDNA density is low and certainly patchy in this vast, heterogenous and dynamic environment. So, the number of sampling replicates and filtered volume necessary to obtain accurate estimates of biodiversity in rich tropical marine environments using eDNA metabarcoding is still unknown. Here, we used a paired sampling design of 30L per replicate on 68 reef transects from 8 sites in three tropical regions and identified fish Molecular Taxonomic Units (MOTUs) using a 12S marker. We quantified local biodiversity variation as MOTU richness, compositional turnover and compositional nestedness between replicated pairs of seawater samples. We report strong turnover of MOTUs between replicated pairs of samples undertaken in the same location, time, and conditions. Paired samples contained non-overlapping assemblages rather than subsets of one-another. As a result, localised diversity accumulation curves showed that even 6 replicates (180L) in the same location underestimated local diversity (for an area <1km). However, sampling of regional diversity using ∼25 replicates in variable locations (often covering 10s of km) achieved saturation of biodiversity accumulation curves. Our results demonstrate high variability of diversity estimates perhaps arising from heterogeneous and local distribution of eDNA distribution in seawater or highly skewed frequencies of eDNA traces. This high compositional variability has consequences for using eDNA to monitor temporal and spatial biodiversity changes of local assemblages. Future biomonitoring efforts could be strongly undermined by a high level of false-negative detections under low replication protocols. We reveal the need to increase replicates or increase sampled water volume to better inform management of marine biodiversity using eDNA.

## Introduction

Biodiversity is changing far faster than our ability to accurately quantify species losses and gains (Ceballos, Ehrlich and Raven, 2020; Filgueiras *et al*., 2021), with consequent difficulties in evaluating the degradation of ecosystem functions and services upon which human well-being depends (Díaz *et al*., 2019). Traditional methods such as visual surveys are very costly, time-consuming and require on-site taxonomic expertise (Kim and Byrne, 2006; Ballesteros-Mejia *et al*., 2013; Dornelas *et al*., 2019). Despite decades of sampling efforts, biodiversity monitoring still covers only a small fraction of global ecosystems and is particularly challenging in isolated and remote regions across the oceans (Collen *et al*., 2009; Webb, Vanden Berghe and O’Dor, 2010; Dornelas *et al*., 2018; Letessier *et al*., 2019). An emerging tool for rapid biodiversity assessment is environmental DNA (eDNA) metabarcoding (Stat *et al*., 2017;Eble *et al*., 2020), which is proving to be particularly effective for marine environments (Juhel *et al*., 2020; Boulanger *et al*., 2021; Holman *et al*., 2021). eDNA-based methods rely on the detection of DNA fragments from various sources including faeces, shed skin cells, organelles, or extruded waste of animals, which become suspended in the water (Dejean *et al*., 2012; Collins *et al*., 2018; Harrison, Sunday and Rogers, 2019). Using filtered water and molecular analyses, eDNA metabarcoding can estimate biodiversity across kingdoms at different taxonomic levels without isolating any target organisms (Valentini *et al*., 2016; Holman *et al*., 2021), and even without exhaustive genetic reference databases (Flynn *et al*., 2015; Juhel *et al*., 2020; Marques *et al*., 2020, 2021). Overall, eDNA metabarcoding has the potential to overcome some limitations of common sampling methods by targeting complete species assemblages, detecting rare (Rees *et al*., 2014), elusive (Boussarie *et al*., 2018) or non-indigenous species (Ficetola *et al*., 2008; Holman *et al*., 2019) and is harmless to organisms and less time-consuming (Bohmann *et al*., 2014; Smart *et al*., 2016).

Yet, routine and widespread eDNA applications to marine ecosystems face multiple challenges (Hansen *et al*., 2018) with variability introduced to biodiversity estimates from multiple sources that are still poorly understood (Bessey *et al*., 2020; Juhel *et al*., 2020; Rourke *et al*., 2021; Thalinger *et al*., 2021). For example, eDNA signals from source-species are weakened by the low biomass-to-water volume ratio and frequent movement of individuals (Moyer *et al*., 2014). Further, long-persistence times, the long-distance transport and patchy aggregation of eDNA by currents, introduce uncertainty linking eDNA signal-to-source (Andruszkiewicz *et al*., 2019; Whitney *et al*., 2021). Detection rates and resultant biodiversity estimates thus critically depend on eDNA *(i) origin* (source of an organism’s genetic material shed into its environment), *(ii) state* (forms of eDNA), *(iii) transport* (e.g. through diffusion, flocculation or settling, currents or biological transport which can vary according to the depth) and *(iv) fate* (how eDNA degrades and decays) (Barnes and Turner, 2016; Harrison, Sunday and Rogers, 2019; Thalinger *et al*., 2021). Besides, water chemistry, salinity and temperature affect the persistence of DNA particles which are best preserved in cold and alkaline waters with low exposure to solar radiation (Moyer *et al*., 2014; Pilliod *et al*., 2014; Strickler, Fremier and Goldberg, 2015; but see Mächler, Osathanunkul and Altermatt, 2018). As a result, marine eDNA residence time ranges from a few hours to a few days and is shorter than in freshwater (Dejean *et al*., 2011; Thomsen *et al*., 2012). In contrast to freshwater systems, marine systems are very open with eDNA particles dispersed by oceanographic dynamics at local (e.g., tides, currents, and water stratification), regional (e.g., eddies) and large (e.g., thermohaline currents) scales in interaction with coastal morphology. Whilst significant dispersal of eDNA from its source may theoretically occur (Andruszkiewicz *et al*., 2019; Eble *et al*., 2020), many studies also indicate that eDNA detection is limited to a small spatio-temporal sampling window even in highly dynamic marine habitats (Port *et al*., 2016; O’Donnell *et al*., 2017; Yamamoto *et al*., 2017; Jeunen, Knapp, Spencer, Lamare, *et al*., 2019; Stat *et al*., 2019; West *et al*., 2020; Boulanger *et al*., 2021). If marine eDNA is sparse, widely transported, and heterogeneously distributed, then biodiversity estimates could be highly variable and eDNA sampling strategies may need to overcome this potentially high noise-to-signal ratio.

The most common approach for concentrating marine eDNA is water filtration along transects (Kumar, Eble and Gaither, 2020) but the appropriate amount of water to filter to overcome sampling variability remains underdetermined (e.g., 1L in Nguyen *et al*., 2020 and 30L in Polanco Fernández *et al*., 2020). An increased volume of water should lead to increased compositional similarly among replicates, but even at 2L 30-50% of the total species pool were missing in any given sample (Bessey *et al*., 2020). The question remains whether a larger water volume, that integrates eDNA signals over multiple kilometres, can provide a less variable and more consistent estimate of biodiversity. If the net effects of ecological and methodological sources of variability are low, we can expect the compositional similarity of repeated 30L eDNA replicates to be high. In contrast, high biodiversity turnover among replicates may suggest further development of sampling protocols is required.

In addition to the volume of water, a high level of eDNA sampling replication in the field can be required to reduce false negatives (species present but not detected) and improve the accuracy of biodiversity estimates. For example, 92 × 2L seawater samples accurately predict (*R*^2^ = 0.92) the distribution of species richness among fish families (Juhel *et al*., 2020). Spatial diversity gradients have been recovered from only three 0.5L water samples in temperate (Thomsen *et al*., 2012) and tropical systems (West *et al*., 2020). However, West *et al*. (2020) report that more replicates were necessary to avoid false-negatives and fully sample diversity in a given site (>8). However, the number of sampling replicates have budget and time limitations (Ficetola *et al*., 2015) – which require optimization to take full advantage of eDNA-based surveys.

Here, we compared biodiversity of replicated eDNA samples in terms of Molecular Operational Taxonomic Units (MOTUs) since genetic reference databases have many gaps for tropical fishes (Marques *et al*., 2020). We assessed within-site MOTU richness, so local or α-diversity, and between-sites MOTU dissimilarity, so *β*-diversity, separating the turnover and nestedness components (Baselga, 2012). We targeted tropical fishes across eight different sites within the Caribbean, Eastern Pacific, and Western Indian Ocean using the same standardised protocol. Over transects 2km long, we filtered 30L of water in a paired sample design. In addition, we performed a replication experiment in two locations by repeating transects multiple times in a ∼24h period. Our objectives were to: (i) establish the extent of variation in fish diversity estimates from replicated eDNA samples collected at the same time, in the same location and under similar conditions, (ii) identify the number of eDNA replicates required to saturate fish diversity at a given site, (iii) compare the above patterns among three ecologically distinct tropical ocean regions, and (iv) examine whether our sampling protocol saturates regional fish biodiversity using between-transect species accumulation curves as an indication of sampling effort appropriate for regional biodiversity estimation. Given that we filtered far more water than previous saturation experiments, we may expect high eDNA recovery rates whereby MOTU richness and composition should be very similar among the paired replicates – providing robust estimates of biodiversity. In this case, the replicate accumulation curve should saturate rapidly and reach an asymptotic maximum. In the opposite case, it would indicate that even a high volume of filtration and a large number of replicates would be required to inventory fish biodiversity regionally.

## Methods

### Sampling sites and eDNA sampling

We filtered surface seawater across eight sampling sites in three different oceanic regions: Caribbean Sea, Western Indian Ocean and Eastern Pacific (Figure 1). At each of the eight sampling sites, several transects were carried out with at least two filtration replicates per transect (see Table 1). Filtration replicates per transect were performed simultaneously on either side of a small boat moving at 2-3 nautical miles per hour while filtering surface seawater for 30 minutes resulting in approximately 30L of water filtered per replicate. The shape of 2km transect varied to match the configuration of the reefs but were always consistent between the compared replicates. eDNA sampling was performed with a filtration system composed of an Athena^®^ peristaltic pump (Proactive Environmental Products LLC, Bradenton, Florida, USA; nominal flow of 1□Lmin^-1^), a VigiDNA® 0.2µM cross flow filtration capsule and disposable sterile tubing for each filtration capsule (SPYGEN, le Bourget du Lac, France). After filtration, the capsules were emptied, filled with 80□mL of CL1 lysis conservation buffer (SPYGEN, le Bourget du Lac, France) and stored at room temperature. A strict contamination protocol in field and laboratory stages was followed using disposable gloves and single-use filtration equipment. More details can be found in Polanco Fernández *et al*. (2020).

**Table 1.** Overview of eDNA sampling across regions and sites in our study. Filtration replicates are our observation units, all samples in column ‘Total filtration replicates’ were used in our accumulation curves. Only paired samples on a given transect were used in the ‘MOTU compositional similarity between filtration replicates’ analyses, indicated by brackets in the ‘Total filtration replicates’ column.

|  | Region | Site | Number of transects per site | Total filtration replicates |
| --- | --- | --- | --- | --- |
| **Local accumulation curves** | Eastern Pacific Ocean | Malpelo | 5 | 10 |
|  | Caribbean Sea | Santa Marta (#1) | 1 | 6 |
|  |  | Santa Marta (#2) | 3 | 6 |
| **Regional accumulation curves** | Western Indian Ocean | Europa | 6 | 12 (12) |
|  |  | Grande Glorieuse | 5 | 10 (10) |
|  |  | Juan de Nova | 5 | 9 (8) |
|  |  | Tromelin | 3 | 6 (6) |
|  | Caribbean Sea | Guadeloupe | 18 | 28 (16) |
|  |  | Providencia | 10 | 20 (20) |
|  | Eastern Pacific Ocean | Malpelo | 13 | 24 (24) |
|  | Caribbean Sea | Santa Marta | 8 | 20 (8) |

**Figure 1.**
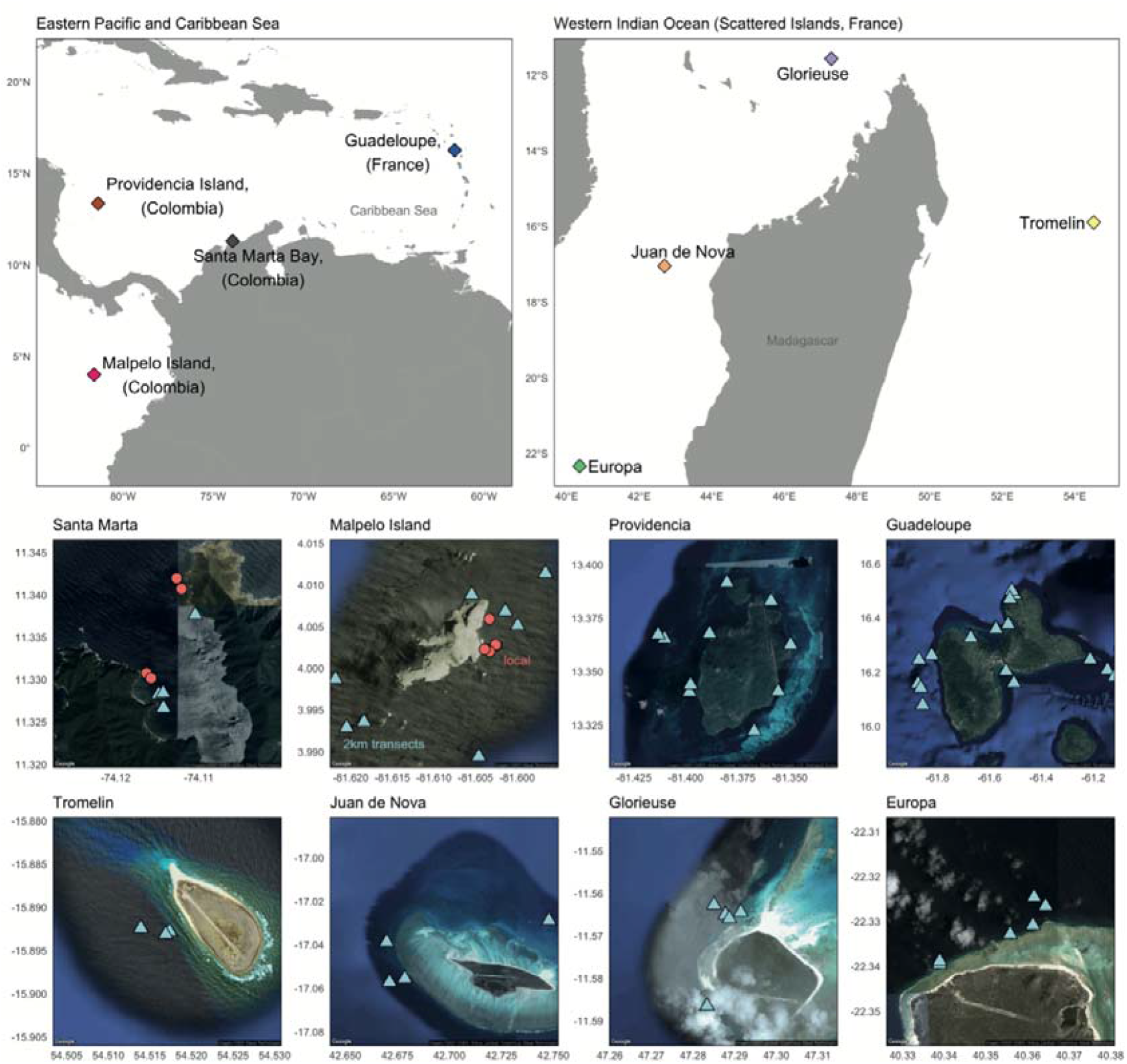
Sampling sites in the Eastern Pacific, Caribbean and Western Indian Ocean. The eight sampled sites represented by Google Earth imagery show the spatial distribution of transects within sites. Markers represent the beginning of eDNA transects in each site; colour and shape indicate whether samples were used in local accumulation analysis (static samples repeated multiple times in a shorter period, red circles) or regional/island level accumulation curves (blue triangles).

### eDNA processing, sequencing, and clustering

eDNA extraction, PCR amplification, and purification prior to library preparation were performed in separate, dedicated rooms following the protocols described in Polanco Fernández *et al*. (2020) and Valentini *et al*. (2016). eDNA was amplified using the teleo primer pair, which targets a marker within the mitochondrial 12S ribosomal RNA gene and shows high accuracy to detect for both bony (*Actinopteri)* and cartilaginous fish *(Chondrichthyes)* (Collins *et al*., 2019). The primers were 5’-labelled with an eight-nucleotide tag unique to each PCR replicate, allowing the assignment of each sequence to the corresponding sample during sequence analysis. Twelve PCR replicates were run per sample, i.e. 24 per transect. Fifteen libraries were prepared using the MetaFast protocol (Fasteris). For seven libraries, paired-end sequencing (2 × 125 bp) was carried out using an Illumina HiSeq 2500 sequencer on a HiSeq Rapid Flow Cell v2 using the HiSeq Rapid SBS Kit v2 (Illumina) and for the remaining eight libraries (Europa, Grande Glorieuse, Juan de Nova, Tromelin) the paired-end sequencing was carried on a MiSeq (2 × 125 bp) with the MiSeq Flow Cell Kit v3 (Illumina), following the manufacturer’s instructions. Library preparation and sequencing were performed at Fasteris facilities. Fifteen negative extraction controls and six negative PCR controls (ultrapure water, 12 replicates) were amplified per primer pair and sequenced in parallel to the samples to monitor possible contaminants.

To provide accurate diversity estimation in the absence of a complete genetic reference database, we used sequence clustering and stringent cleaning thresholds (Marques et al., 2020). Clustering was performed using the SWARM algorithm which uses sequence similarity and abundance patterns to cluster multiple variants of sequences into MOTUs (Fisher *et al*., 2015b; Rognes *et al*., 2016). First, sequences were merged using *vsearch* (Rognes *et al*., 2016), cutadapt (Martin, 2011) was then used for demultiplexing and primer trimming and *vsearch* to remove sequences containing ambiguities. SWARM was then run with a minimum distance of one mismatch to make clusters (Marques et al., 2020). Once the MOTUs are generated, the most abundant sequence within each cluster is used as a representative sequence for taxonomic assignment. Then, a post-clustering curation algorithm (LULU) was applied to curate the data (Frøslev *et al*., 2017). We finally removed all occurrences with less than 10 reads per PCR and all MOTUs present in only one PCR within the entire dataset. This additional step is necessary as PCR errors are unlikely to be present in more than one PCR occurrence and it removes spurious MOTUs inflating diversity estimates; see Marques *et al*., (2020) for more details.

### MOTU richness

We first compared MOTU local richness with the expected richness of the species pool in the eight sites. For this, we created a MOTU presence-absence matrix containing every replicate of each region and compiled fish species lists for each of the eight sites from the literature: Scattered Islands (Grande Glorieuse, number of species = 576; Europa Island, n = 506; Juan de Nova Island, n = 480; Tromelin Island, n = 239; personal communication with Terres Australes et Antarctiques Francais; www.taaf.fr), Tayrona Park (n = 515; SIBM, 2021), Providencia (n = 343; Robertson and Van Tassell, 2019), Malpelo (n = 257; Robertson and Allen, 2015) and Guadeloupe (n = 425; Froese and Pauly, 2000). To examine whether the transect MOTU richness varied among oceanic regions (n=3), we performed a Kruskal-Wallis rank sum test and related the MOTU richness per replicate to the site richness (from species lists) using a linear model. We also estimated the recovered MOTU richness for each filtration replicate per transect. We determined if the mean α-diversity differed between paired filtration replicates for a given transect using a Wilcoxon signed rank test.

### MOTU compositional dissimilarity

To understand the variability in MOTUs recovered between filtration replicates we quantified the compositional similarity of MOTUs. We estimated the pairwise Jaccard’s dissimilarity index (β_jac_) between filtration replicates per transect using the R package *vegan* (Oksanen *et al*., 2019). The Jaccard index ranges from 0 (species composition between the replicates is identical, i.e., complete similarity) to 1 (no species in common between the replicates, i.e., complete dissimilarity). We partitioned the Jaccard index into turnover (β_jtu_) and nestedness (β_jne_) components using the R package *betapart* (Baselga and Orme, 2012). Nestedness quantifies the extent to which replicates are subsets of each other. Turnover indicates the amount of species replacement among replicates, i.e., the substitution of species in one replicate by different species in the other one (Baselga and Orme, 2012; Legendre and De Cáceres, 2013). In addition, we tested whether β_jac_ differed between the regions using a Kruskal-Wallis rank sum test.

### Local-scale MOTU accumulation curves

To analyse the local-scale richness accumulation, we repeated circular transects multiple times in Malpelo and Santa Marta. We sampled two locations in Santa Marta filtrating 6 replicates at each within 20 hours, and one location in Malpelo filtrating 10 replicates within three days. This sampling design defined three local MOTU accumulation ‘experiments’. We produced MOTU richness accumulation curves across filtration replicates from each location using the *specaccum* function from the R package *vegan* (Oksanen *et al*., 2019). The ‘*random’* method was used to generate 1,000 accumulation curves, which were used to fit models describing the relationship between the number of replicates and MOTU richness. We fitted fourteen models to each saturation experiment and ranked fitted models by AIC score. We generated multi-model mean averages which were used for asymptote calculations, extrapolation and visualisation using the *sars_average* function from the *sars* R package (Matthews *et al*., 2019). We next used the *sar_pred* function to extrapolate MOTU richness for up to 60 filtration replicates. We defined asymptotes as the number of replicates at which less than 1 new MOTU was added per additional sample.

### Regional-scale MOTU accumulation curves

In contrast to the saturation curves at one location, we assessed the extent to which our eDNA protocol captures regional fish biodiversity. MOTU accumulation curves were calculated using all filtration replicates in each of the 8 sites. Species accumulation curves were produced and compared as above (Figure 1; Table 1) rather than within localised repeated transects. All transects and replicates from all stations within a sampling site were pooled to form a site-wide (or regional) accumulation curve.

All analyses were performed in R version 4.0.1 (R Core Team, 2020). The activities in Malpelo were undertaken with the permit: “Resolución Número 0170-2018 MD-DIMAR-SUBDEMAR-ALIT 8 de marzo de 2018”.

## Results

### Overview of eDNA biodiversity patterns

We detected a total of 789 unique MOTUs assigned to bony and cartilaginous fish taxa. Site MOTU richness was significantly and positively associated with the size of the site species pool (slope=0.1, t=4.7, p<0.001; Figure 2) reconstructing large-scale biodiversity gradients across the tropics.

**Figure 2.**
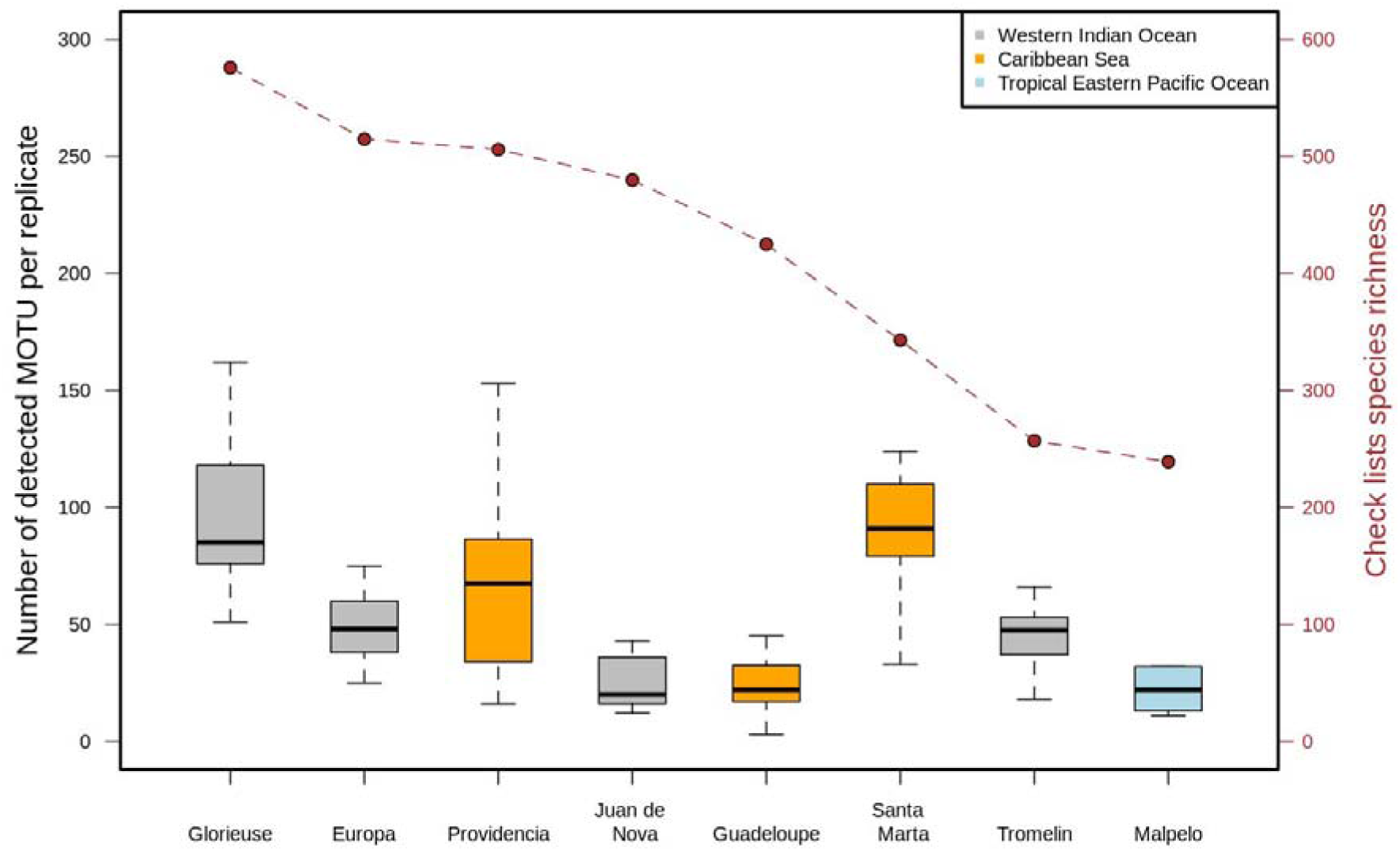
Sites ranked by their expected species richness based on species lists in different regions (see Methods). One single filtration replicate recovered up to 162 fish MOTUs in the most diverse site monitored. Overall, we found an association between the size of the species pool and the number of MOTUs recovered in a single replicate. The relatively consistent proportional MOTU samples from the full species pools suggests that in richer sites filtration replicates were not saturated with available eDNA. The bold central lines correspond to median values across transects (filtration replicate pairs), interquartile range (25th-75th) corresponds to box edges, and whiskers extend to 1.5 x interquartile range.

### MOTUs richness per replicate

The fish MOTU richness detected by each filtration replicate (n = 100) ranged from 3 to 162, with a mean of 58.3 ± 35.6 MOTUs (Figure 2). The mean α-diversity detected by each filtration replicate for a given transect did not differ significantly (Wilcoxon signed rank test: n =50, Z = -0.927, p = 0.354). The MOTU richness detected at each transect (i.e., two filtration replicates combined, n = 50) ranged from 19 to 184, with a mean of 82.5 ± 42.7 MOTUs and did not differ significantly among regions (Kruskal-Wallis rank sum test: χ^2^ = 4.0682; p = 0.1308). On average, 69.7% of the MOTU richness along a transect was identified by a single filtration replicate, ranging from 11.5% to 98.1%, with variations among regions (Western Indian Ocean = 63.6%, n = 36; Eastern Pacific = 74.5%, n = 20; Caribbean = 72.5%, n = 44).

### MOTU compositional dissimilarity between replicates

The composition of fish MOTUs was highly dissimilar between paired replicates (Figure 3; mean similarity = 0.598 ± 0.155 where 1 is full dissimilarity with no MOTU is common), and varied among transects ranging from 0.174 to 0.882. The level of dissimilarity between paired replicates varied significantly among regions (Kruskal-Wallis rank sum test: χ^2^ = 22.791; p < 0.001) being most dissimilar in the West Indian Ocean (mean=0.729 ± 0.102) than in the Caribbean (0.528 ± 0.146) and the Eastern Pacific (0.511 ± 0.081). MOTU compositional differences between replicates were primarily due to MOTU turnover (Figure 3; n = 49, Z = -6.097, p < 0.001; mean turnover = 0.450 ± 0.153) with a lower contribution of nestedness (mean = 0.149 ± 0.146). Turnover ranged from 0.095 to 0.846 and nestedness from 0.005 to 0.646.

**Figure 3.**
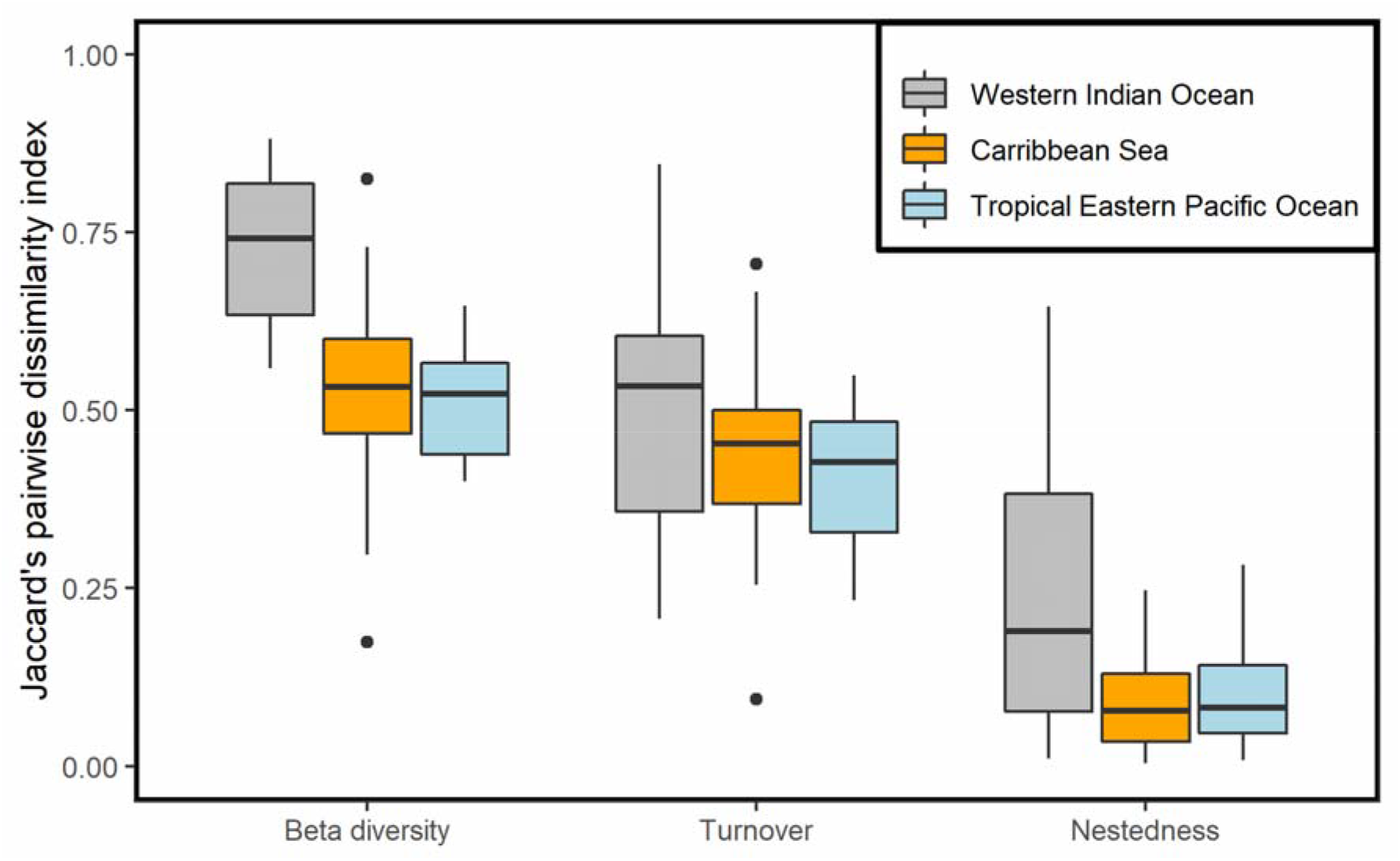
Boxplots of the MOTU compositional dissimilarity between eDNA filtration replicates (β-diversity), and deconstruction into turnover and nestedness components for three ocean regions. The bold central lines correspond to median values across transects (filtration replicate pairs), interquartile range (25th-75th) corresponds to box edges, and whiskers extend to 1.5 x interquartile range.

### Local-scale MOTU accumulation curves

The accumulated fish MOTU richness in the two locations in Santa Marta was between 109 and 131 and the one location was 114. After 6-10 replicates sampling in the same location, MOTU richness did not fully saturate with additional replicates adding new MOTUs to the total (Figure 4). Modelled accumulation curves suggest that 27-58 filtration replicates would be required to reach an asymptotic richness of 164-251 MOTUs in Santa Marta, and 23 filtration replicates to reach an asymptotic richness of 134 MOTUs in Malpelo.

**Figure 4.**
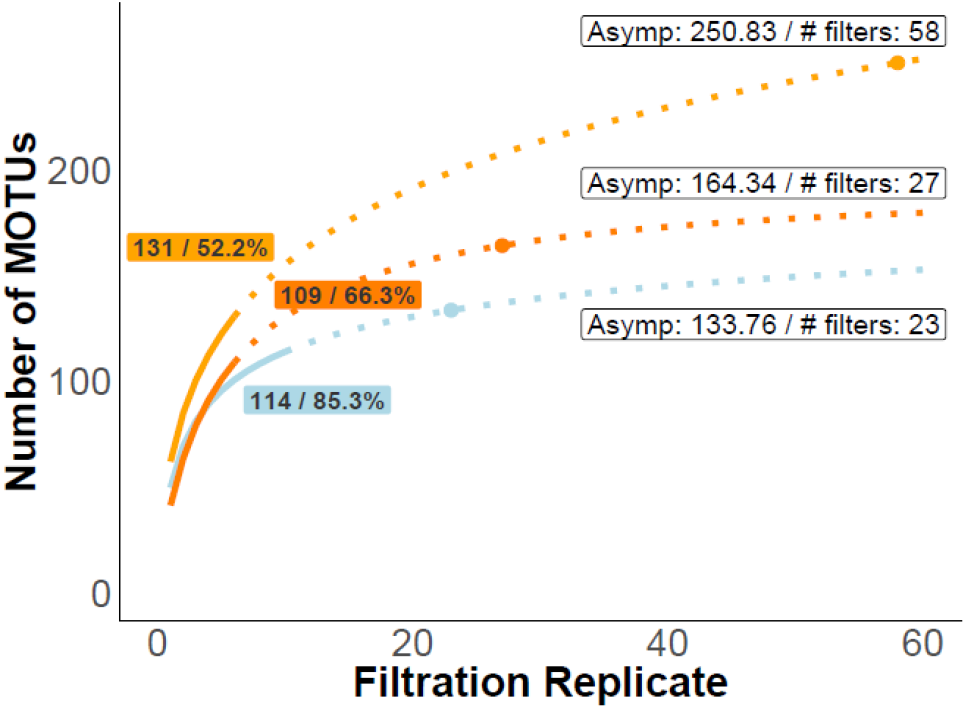
Local-scale MOTU richness accumulation analysis of eDNA filtration replicates from Santa Marta and Malpelo. The curves show the multi-model mean average of the local MOTU richness and richness extrapolation for the filtration replicates collected by repeated sampling at the same location over a short period. Coloured text boxes indicate the final sampled richness and the percentage of the estimated richness asymptote reached with our filtration replicates. Points on the curve mark the asymptote (defined as a < 1 MOTU increase in species richness per added sample). The asymptotic MOTU richness plus the number of filters required to reach the asymptote are noted in the white text box next to the curves. The solid line shows the richness of the filters collected during actual sampling; the dotted line is the extrapolation of richness up to 60 filters. The curve colour corresponds to the sampling regions: Santa Marta (light orange: ‘tayrona_camera_1’, dark orange: ‘tayrona_camera_2’), Malpelo (blue).

### Regional-scale MOTU accumulation curves

At a regional scale, MOTU accumulation curves detected various proportions of the total asymptotic MOTU richness (defined as <1 additional MOTU per filtration replicate; 98.8% on average in the Caribbean Sea, 103.3% on average in Malpelo, 67.4% on average in the Western Indian Ocean). The Caribbean and Eastern Pacific filtration replicates saturated MOTU richness after 18 to 28 replicates (i.e., within our number of replicates), except for Santa Marta, where an additional 6 replicates are predicted to be required to reach an asymptote (Figure 5). In the Western Indian Ocean, where sampling was less exhaustive, regional MOTU richness did not saturate and reached between 46.4% (Tromelin) and 82.7% (Grande Glorieuse) of the predicted asymptotic MOTU richness. To reach an asymptotic richness of 172.3-320.2 MOTUs in the Western Indian Ocean, our estimates suggest that between 30-52 additional replicates would be required. The shapes of regional accumulation curves were qualitatively different between the three oceans and showed differing levels of both diversity and sampling exhaustiveness across sites (Figure 5).

**Figure 5.**
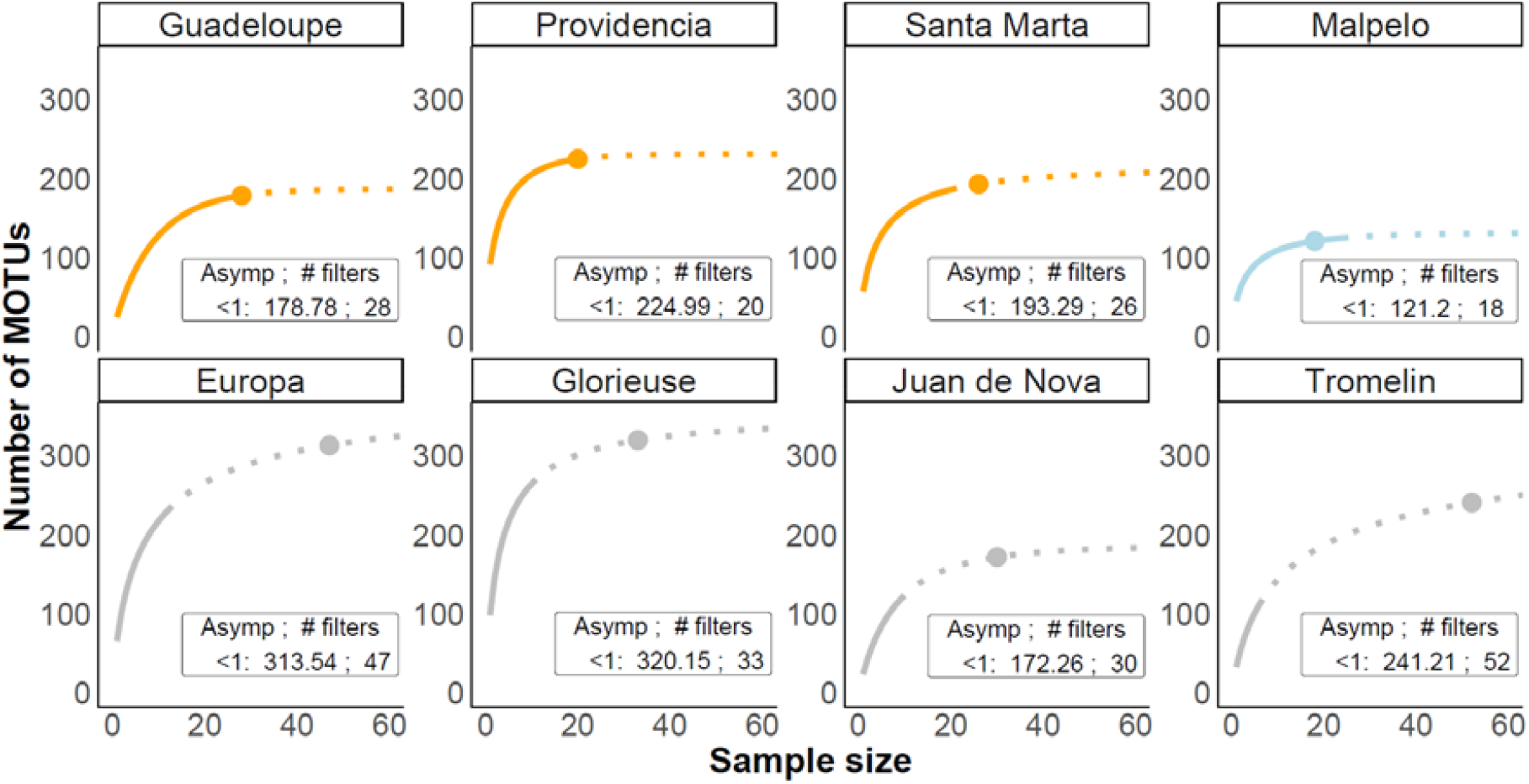
Regional MOTU richness accumulation curves of eDNA filtration replicates across the Caribbean, Eastern Pacific and Western Indian Ocean. The curves show the multi-model mean averages of the local richness and richness extrapolation (number of MOTUs) for the number of filters (sample size) from each region. Points on the curve represent the asymptote (defined as a less than 1 MOTU increase in species richness per added sample). The asymptote for the MOTU richness plus the number of filters needed to reach the asymptote are noted in the text box below the curves. The solid line shows the richness of the filters collected; the dotted line is the extrapolation of richness up to 60 filters. The colours of the curves correspond to the sampling area: Caribbean Sea (orange), Eastern Pacific (light blue), Western Indian Ocean (grey).

## Discussion

Our results reveal that, without adequate replication, sampling variability undermines biodiversity estimates in highly diverse tropical ecosystems, such as coral reefs (Cilleros *et al*., 2019; Bessey *et al*., 2020; Juhel *et al*., 2020; Polanco Fernández *et al*., 2020). The variability of eDNA biodiversity estimates showed high compositional dissimilarity between filtration replicates so, for some locations, even extensive sampling with 6-20x 30L replicates could reach an asymptotic number of detected MOTUs. Promisingly, there was little difference in overall MOTU richness between paired replicates, although these replicates recovered different species identities so are essential in eDNA sampling design.

The similar MOTU richness but different composition between replicates suggest that eDNA distribution varies at a fine-scale in seawater and is certainly patchier than previously thought. This patchiness of fish eDNA in tropical reefs is further supported by Bessey *et al*. (2020) who report that multiple collections from a single site 2m apart have <30% overlapping species detections. The fine-scale distribution of eDNA in the environment could be a function of multiple factors. For example, ambient eDNA in seawater could be modified by complex sea currents, surface slicks (Whitney *et al*., 2021), local water dynamics, thermohaline circulation forces, spatio-temporal variation in organism activity and behaviours (e.g., spawning, feeding, diel migrations), different DNA shedding, degradation and decay rates (Harrison, Sunday and Rogers, 2019). Identifying when, where, and to what extent these varying processes act to modify spatial and temporal eDNA distribution is critical to disentangle biodiversity variation from sampling variation on reefs.

Since biodiversity changes are most often detected as compositional turnover, but not necessarily richness changes, we highlight a major challenge in developing eDNA to monitor ecosystem modifications through space and time (Dornelas *et al*., 2014; Hill *et al*., 2016; Santini *et al*., 2017; Blowes *et al*., 2019). Our results imply that if sample variability is not accounted for, or survey designs are not well replicated, eDNA-derived time-series could over-emphasise compositional turnover by containing many false-negatives. This point will be exacerbated where incomplete reference databases recover a small portion of common species and falsely identify low species turnover among samples (Schenekar *et al*., 2020), even though MOTU turnover identified here may be very high. We found MOTU compositional differences between replicates to be higher in the more speciose Western Indian Ocean (under similar sampling protocols), perhaps due to the larger species pool, further challenging eDNA applications in most diverse tropical systems (Juhel et al. 2020). Current protocols should be cautiously applied to biomonitoring if such limitations remain unresolved. Our results also imply that many replicates of >30L water are needed to reach a stable estimate of total local biodiversity. Promisingly, the regional biodiversity of tropical systems was relatively well-quantified through repeated eDNA sampling (e.g., Figure 5), and more exhaustive biodiversity estimates may be achieved by including mesophotic coral ecosystems (from -300m depth to sub-surface) and various habitats (e.g. lagoons, reef-slope, mangroves, sea grass) (Juhel *et al*. 2020).

The well-established community pattern that many species are rare and few are common (McGill *et al*., 2007) also likely exists in eDNA particles. Moreover, finding rare eDNA fragments in any given sample may be exacerbated by features of marine systems. For example, we likely sampled vagrant open-ocean species, which only pass through temporarily, in some of our remote sites (e.g., Malpelo) which may have increased sampling variability. Compared to terrestrial systems, the seawater environment may homogenise eDNA that comes from different habitats (e.g., coral, rock, sand, seagrass). The eDNA species pool could be larger in a seawater sample than expected based on habitat variation along a given transect. Dispersion of eDNA between distinct habitats (e.g., from seagrass beds to coral reefs) would enhance the likelihood of finding a rare habitat specialist from a different habitat type and increasing perceived sampling variability. As such, eDNA variability may be greater in seascapes with a greater diversity of habitats. Sampling designs may need to account for the extent that a given water body accumulates sources of eDNA, and the amount of habitat variation that a water sample signal is aggregated over. eDNA statistical analyses may also need to control for habitat variations before reaching conclusions (Boulanger *et al*., 2021).

Marine eDNA protocols are challenged by the compositional turnover between replicates. As in traditional approaches, saturation of biodiversity samples only occurs with many replicates on tropical reefs (MacNeil *et al*., 2008). However, traditional methods like underwater visual census (UVC) and baited remote underwater video (BRUVs) are systematically biased by observer effects and fish behaviour, leading to false negatives for cryptic and elusive species (Ackerman and Bellwood, 2000; MacNeil *et al*., 2008; Bernard *et al*., 2013). For example, we found ∼30 Chondrichthyes species that typically would not be encountered on visual surveys (e.g., 2 *Mobula sp*., 6 *Carcharhinus sp*.; Polanco Fernández *et al*., 2020), among other elusive and endangered megafauna that have been uncovered during similar sampling regimes (Juhel *et al*., 2021). We highlight that eDNA replicates may be affected by factors that contribute to the precision of biodiversity estimates, rather than a biased biodiversity signal as obtained with UVC or BRUVs, e.g. a lack of eDNA in water samples leading to availability errors; although see Stat *et al*., (2019) for biases against specific genera and Kelly, Shelton and Gallego, (2019) for discussions of primer efficiency biases.

Coral reefs are extremely speciose (Fisher *et al*., 2015a; Edgar *et al*., 2017), and thus 60L (two replicates) or even 180L (six replicates) does not seem to capture the extent of local biodiversity. Instead, in support of other eDNA studies that filtered far less water, our replicates only sampled a portion of diversity (DiBattista *et al*., 2017; Jeunen, Knapp, Spencer, Taylor, *et al*., 2019; Koziol *et al*., 2019; Sigsgaard *et al*., 2019; Stat *et al*., 2019; Bessey *et al*., 2020; Juhel *et al*., 2020). In temperate systems, 20L of water was sufficient for fish family richness to saturate (Koziol *et al*., 2019; but see Evans *et al*., 2017), but tropical systems are more challenging to monitor. The number of eDNA replicates to ensure tropical fish diversity saturation varies widely. For example, 32-39 samples of 0.5L of water began to saturate fish genera diversity in western Australia (Stat *et al*., 2019), but 92 samples of 2L did not saturate diversity in West Papua, Indonesia, a hotspot of fish diversity (Juhel *et al*., 2020). Furthermore, even the largest sample of 2L in Bessey *et al*. (2020) only detected <43% (75/176) of the total species pool reported in the Timor Sea.

eDNA accumulation curves often confound site-accumulated (regional) and replicate-accumulated (local) diversity presenting challenges for replicate number and water volume refinements (but see Bessey *et al*. 2020). Comparing available estimates, integrative sampling (performed here), rather than point sampling, e.g., Stat *et al*., (2019) and Juhel *et al*. (2020), appears very promising. For example, in Caribbean and Eastern Pacific sites within ∼25 filters we find additional filters added only <1 MOTU. Previous works using point samples have far higher sampling numbers, and higher bioinformatic costs per filter so leading to apparently lower cost-effectiveness (unless filters are aggregated at the DNA extraction step; e.g., Stat *et al*., 2019; Juhel *et al*., 2020). Future work should optimise sampling designs and the trade-off between water sample volume and replicate number, which we only partially explore, and how these factors contribute to the precision of biodiversity estimates in controlled settings (Miya *et al*., 2015). For example, if sampling nearer to substrate bottoms greatly improves recovery of eDNA this additional cost (e.g., divers, submersibles, and additional expertise) could work out as a cost-effective solution to address surface sampling variability. Another option would be to use previous knowledge of biodiversity in each site to adapt the number of replicates to reach expected saturation.

A similar pattern of low compositional similarity, and consistent richness in replicates, could arise if filters saturate with eDNA and prevent the full quantification of biodiversity. Our analyses suggest this is unlikely because the richness recovered from the eDNA filters was associated with the size of the species pools, which would be unexpected if filters had a maximum richness capacity that was reached consistently. Furthermore, we might expect nestedness to be more important if filters or PCR processes were first saturated with the most commonly available eDNA, but we found MOTU compositional differences between replicates were more strongly related to turnover than nestedness. Finally, if filters first saturate with common species, eDNA recovery of rare species would be limited but in our eDNA protocol we find many species that remain undetected or rare in visual surveys (Polanco Fernández *et al*., 2020). Promisingly, this suggests not only that our sampling protocol is robust but also that sampling and filtering an even greater water volume per filtration replicate is a feasible approach to better quantify the high fish diversity of coral reefs. Given the low biomass-to-water ratio in marine systems a high volume of filtered water is likely a prerequisite to have a representative sampling of the marine environment (Bessey *et al*., 2020). However, other parameters must be considered and explored in the future to identify whether physicochemical and local oceanographic conditions introduce variability in biodiversity estimates.

## Conclusion

Our findings underline both the promises and limitations of eDNA derived biodiversity estimates in hyper-diverse tropical ecosystems. On one hand, local richness estimation appears to rapidly resolve broad-scale richness patterns of under-documented tropical marine biodiversity (Costello *et al*., 2010; Menegotto and Rangel, 2018). On the other hand, high stochasticity between samples urges cautious application to biomonitoring, and further protocol refinement, to avoid misattribution of biodiversity trends to detection errors. A better understanding of the behaviour of eDNA in diverse physicochemical marine environments will help design more effective eDNA sampling protocols and disentangle sampling errors from true biodiversity patterns (Harrison, Sunday and Rogers, 2019). Resolving whether more replicates, or greater water volumes, leads to higher probability of eDNA recovery is critical for cost-effective eDNA protocols – but integrative sampling of tens of litres along boat transects appears a promising approach. We also recommend testing various water sampling strategies, for example sampling not only surface water, but taking eDNA along a depth gradient where the ecology of eDNA may differ. Physicochemical parameters of the water bodies could be important to consider when designing the eDNA sampling strategy. Accurate, cheap and fast biodiversity estimates are critically needed to monitor changes in the Anthropic ocean. Current eDNA protocols provide higher and more realistic estimates of biodiversity than traditional methods for a given sampling effort. This opens very promising and realistic perspectives to quantify biodiversity since increasing the volume of water filtered and replicates numbers is feasible, particularly in regions with high biodiversity. Further refinement of our marine eDNA protocol will better quantify, monitor, and manage changing tropical marine biodiversity.

## Acknowledgements

This project was supported by the ETH Global grant and the Monaco Explorations Foundation, CORALINA, which provided support for entrance to the island of Providencia for the development of the project and maritime support for the field sampling. SB and FL were supported by the Fondo Patrimonial de Malpelo, Parques Nacionales Naturales de Colombia and Ministerio de Ambiente de Colombia. GHBP, MMM and APF were supported by the Instituto de Investigaciones Marinas y Costeras (INVEMAR) through the project “Investigación científica hacia la generación de información y conocimiento de las zonas marinas y costeras de interés de la nación”, BPIN code 2017011000113. LP received funding from the Swiss National Science Foundation for the project Reefish (grant number 310030E-164294). CA was funded by an “étoile montante’’ fellowship from the “pays de la loire” region (grant number 2020_10792). EM was supported by the FAIRFISH project (ERC starting grant: 759457). We thank SPYGEN staff for technical support in the laboratory and are grateful to PE Guerin for his support in bioinformatics pipeline development.

## Data Accessibility Statement

We agree to archive our data in a public repository on acceptance.

